# Metabolic predictors of response to immune checkpoint blockade therapy

**DOI:** 10.1101/2023.04.21.537496

**Authors:** Ofir Shorer, Keren Yizhak

## Abstract

Metabolism of immune cells in the tumor microenvironment (TME) plays a critical role in cancer patient response to immune checkpoint inhibitors (ICI). Yet, a metabolic characterization of immune cells in the TME of patients treated with ICI is lacking. To bridge this gap we performed a semi-supervised analysis of ∼1700 metabolic genes using single-cell RNA-seq data of >1 million immune cells from ∼230 tumor and blood samples treated with ICI. When clustering cells based on their metabolic gene expression, we found that similar immunological states are found in different metabolic states. Most importantly, we found metabolic states that are significantly associated with patient response. We then built a metabolic predictor based on a dozen gene signature which significantly differentiates between responding and non-responding patients across different cancer types (AUC = 0.8-0.86). Taken together, our results demonstrate the importance of metabolism in predicting patient response to ICI.

## Introduction

Over the past decade cancer therapy has been revolutionized by the usage of immune checkpoint inhibitors (ICI), resulting in unprecedented rates of long-lasting tumor responses in patients with a variety of cancers^1^. Nevertheless, most patients do not respond to treatment or acquire resistance^2–4^, reaching response rates of less than 40% in solid tumors^5^ with a large number of partial responders^6^. To address this challenge, multiple biomarkers of response have been suggested, including different gene signatures^4,7,8^, T-cell abundance levels^9–11^, as well as metrics related to the tumor’s genetics, including the tumor mutational burden and copy number alteration^12–16^. These and other metrics are however rarely predictive across datasets, and certainly across different cancer types.

More recently, differences in the metabolic adaptation of tumor and immune cells and their competition for resources have been shown to contribute to patient response^17–20^. Changes in cellular metabolism result with tumor-immune interactions in a way that both deplete essential nutrients from the shared environment and generate immunosuppressive metabolites^20,21^. For instance, enzymatic activation of IDO depletes tryptophan levels, a vital nutrient for T cell response. At the same time, this activity generates kynurenine, a metabolite causing upregulation of PD1 on activated CD8^+^ T cells^22,23^. Another example involves the high lactate levels generated by tumor cells due to up-regulation of LDH, resulting in an acidic environment that suppresses NK cells cytotoxicity, blocks monocytes and dendritic cells differentiation, and inhibits effector T cells activity^24–27^. These and other findings were mainly established in *in vitro* systems or in animal models. Studying the metabolic transcriptomic changes that occur in immune cells found in the TME of patients treated with ICI, can therefore shed light on mechanisms of response to therapy and highlight immune metabolic vulnerabilities.

Here we perform a semi-supervised analysis of single-cell RNA-seq data from tumor biopsies of patients treated with ICI, focusing on ∼1700 metabolic genes that are associated with ∼100 metabolic pathways^28^. Focusing on this well-defined cellular process and its narrowed gene set, enabled us to discover novel associations with patient response that were masked under the genome-wide analyses. We found that different immune cell states share similar metabolic activity and vice versa, resulting in a novel classification to cellular states, mainly involving CD8^+^ T-cells. Furthermore, we detected metabolic clusters that are significantly associated with patient response. A non-discrete analysis of metabolic programs revealed a metabolic gene signature that is significantly associated with tumor progression or regression across various cancer types, including melanoma, Merkel cell carcinoma, lung, and breast cancers. Finally, we show a metabolic involvement in the polarization of macrophages to a suppressive M2-like phenotype in an acquired resistance patient, together with the associated tumor metabolic changes.

## Results

### Immune cellular metabolism and its association with patient response

To identify immune metabolic changes at the single-cell level and study their association with ICI treatment and patient response, we re-analyzed our previously published dataset that includes single-cell RNA sequencing (scRNA-seq) of 16,291 CD45^+^ cells from 48 tumor biopsies taken from 32 metastatic melanoma patients treated with ICI^29^. 11 patients had longitudinal biopsies and 20 patients with one or two biopsies, taken either at baseline or during treatment. Tumor samples were classified based on radiologic assessments into progression/non-responder (NR; n = 31, including SD/PD samples) or regression/responder (R; n = 17, including CR/PR samples)^30^. Using the original cell classification into 11 clusters, we devised a score for 97 metabolic pathways based on the expression of their associated metabolic genes^28^ (Supplementary Table S2, Methods).

This analysis showed several metabolic pathways which are significantly more expressed than others across all cells, as well as specific cell types and states that are more metabolically active (Figure 1b, c, Supplementary Figure S1). Specifically, we found that pathways such as glycolysis, oxidative phosphorylation (OXPHOS) and tricarboxylic acid (TCA) cycle, are generally active across many cell types, as was similarly demonstrated in melanoma and head and neck squamous cell carcinoma (HNSCC) patients^31^. In contrast, other pathways such as nucleotide salvage pathway, heme degradation, and purine synthesis, are active only in specific cell types (Figure 1d). Moreover, we found that cells from the myeloid lineage and cycling exhausted T cells, that were previously associated with negative response to immunotherapy^29^, are the most metabolically active. On the other hand, memory T cells and B cells that were previously associated with positive response to immunotherapy^29^, show lower metabolic expression across all pathways (Figure 1c).

**Fig. 1.**
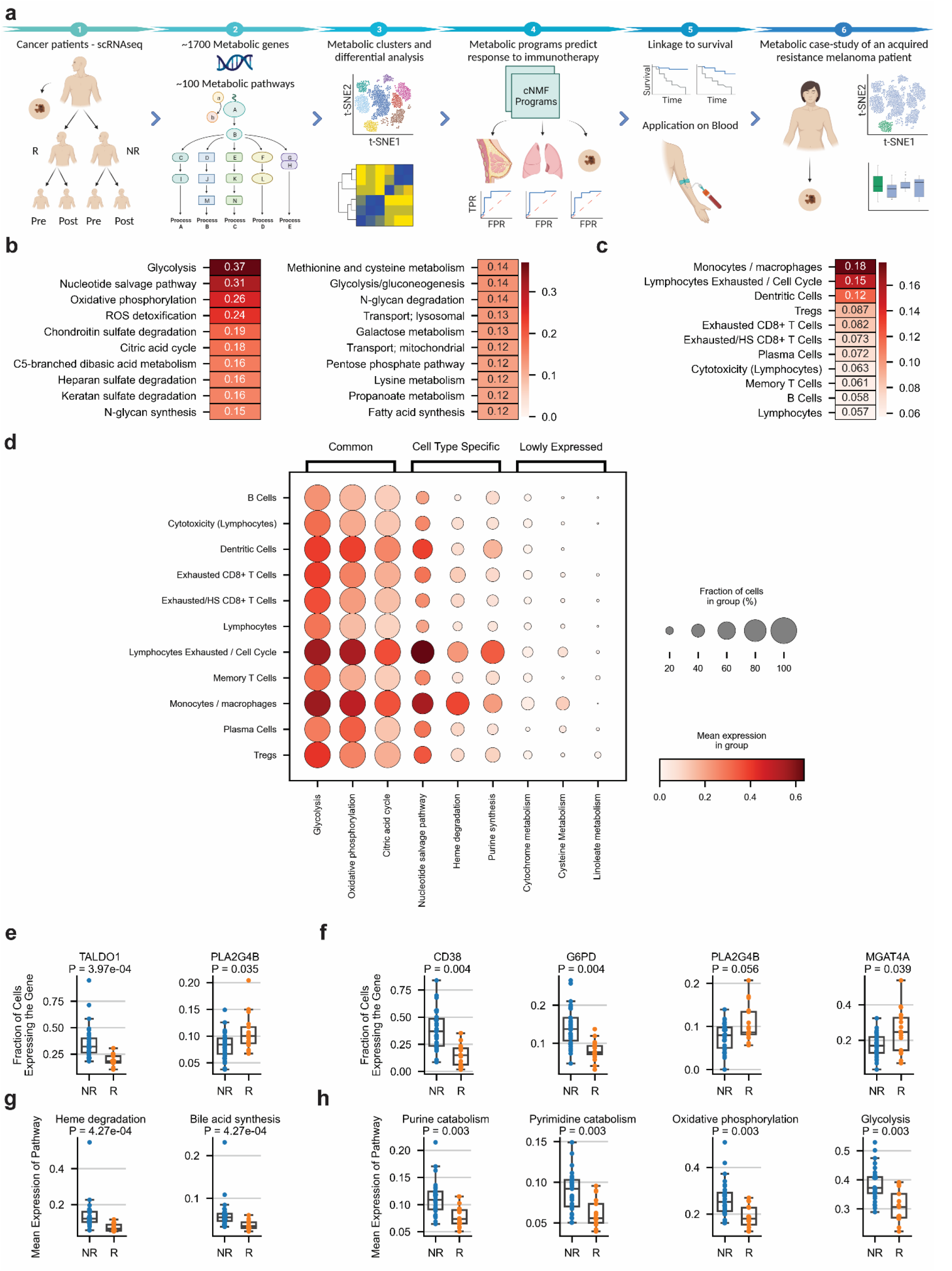
Immune cellular metabolism and its association with patient response. a. Schematic workflow of the study. b. Top 20 active metabolic pathways across all single-cells. c. Rank of metabolic activity by immune cell types. d. Selected metabolic pathways across different cell types separated by commonly expressed, cell-type specific, and lowly expressed pathways. e. Fraction of cells expressing metabolic genes associated with patient response across all cells. f. Fraction of cells expressing metabolic genes associated with patient response across CD8^+^ T Cells. g. Mean expression of metabolic pathways associated with non-responding samples across all cells. h. Mean expression of metabolic pathways associated with non-responding samples across CD8^+^ T Cells. Abbreviations: R = Responders, NR = Non-responders.

Examining the association between metabolic genes and patient response to ICI, we found a couple of genes that are significantly up-regulated in responding samples (PLA2G4B & DGKD), both are involved in Glycerophospholipid metabolism (Figure 1e, Supplementary Table S3). This pathway has shown to be involved in different immune cellular functions, including T cell proliferation and activation, and macrophages differentiation^32–34^. Additional genes were found to be significantly up-regulated in non-responding samples, including glycolytic genes (PGAM1, GPI, TPI1, ENO1, GAPDH), pentose phosphate pathway genes (TALDO1, G6PD), TCA cycle genes (IDH2, MDH2), and oxidative phosphorylation genes (NDUFA9, NDUFA12, UQCRFS1, NDUFA13, NDUFB4, COX5B, Figure 1e, Supplementary Table S3). When focusing only on CD8^+^ T cells, the N-glycosylation enzyme MGAT4A and PLA2G4B were significantly expressed in responders, while G6PD, CD38, MDH2, DBI and others, were significantly expressed in non-responders (Figure 1f, Supplementary Table S3). At the pathway level, while no metabolic pathways were found to be significantly expressed in responding samples (Supplementary Table S3), in non-responding samples we saw a significant expression of pathways such as heme degradation, bile acid synthesis, purine catabolism, oxidative phosphorylation, fatty acid oxidation, and glycolysis (Figure 1g, Supplementary Table S3). Repeating the pathway analysis using only CD8^+^ T Cells, we found a significant expression of multiple pathways including purine catabolism, glycolysis, OXPHOS, and pyrimidine catabolism in non-responding samples (Figure 1h, Supplementary Table S3). Of note, as previously found in the genome-wide analysis^29^, no significant metabolic changes were found between baseline and post-treatment samples. Overall, our analysis identified specific cell states that are more metabolically active, and metabolic gene markers associated with regression or progression of individual tumors in response to ICI therapy.

### Metabolic-based clusters of melanoma infiltrated immune cells and their association with patient response to ICI

To metabolically define the different immune cells in an unbiased manner, we clustered all ∼16,000 cells using the pre-defined set of 1689 metabolic genes^28^ (Supplementary Table S2). This process yielded 10 distinct metabolic clusters that are partially different from the 11 original clusters obtained using all genes (Figure 2a, Figure 2b). Specifically, the previously defined clusters of B- and plasma-cells (C5 and C9), cycling cells (C6) and myeloid cells (C7, C8 and C10), were also found to have a distinct metabolic signature (Figure 2b, Supplementary Table S3, Supplementary Figure S2): C5 significantly expressed genes related to lipid and glycan metabolism, such as PLCG2, ALOX5, PIK3C2B, ST6GAL1 and MGAT5; C6 was enriched with genes of OXPHOS, TCA and fatty acid oxidation (FAO), as well as genes related to nucleotide interconversion (RRM1, RRM2, DUT, TK1, TYMS) and genes involved in folate metabolism (DHFR, GGH, MTHFD1 & FPGS), all contribute to cell proliferation. Macrophages were divided into two metabolic clusters (C7 and C8), where C8 demonstrated a general upregulation of metabolic pathways compared to C7, including OXPHOS, TCA cycle, metabolism of fructose and mannose, pyrimidine synthesis, cholesterol metabolism, and phosphatidylinositol phosphate metabolism (Supplementary Table S3). When comparing the two clusters at the gene level, C8 significantly expressed genes related to its enriched metabolic pathways, including apolipoproteins (APOC1, APOC2, APOE), OXPHOS genes (NDUF genes, COX genes & CYC1), and TCA cycle genes (IDH1, MDH1, SDHA, SDHB, SDHD & FH), while C7 expressed more IDO1, AGPAT9 and SDS (Supplementary Table S3).

**Fig. 2.**
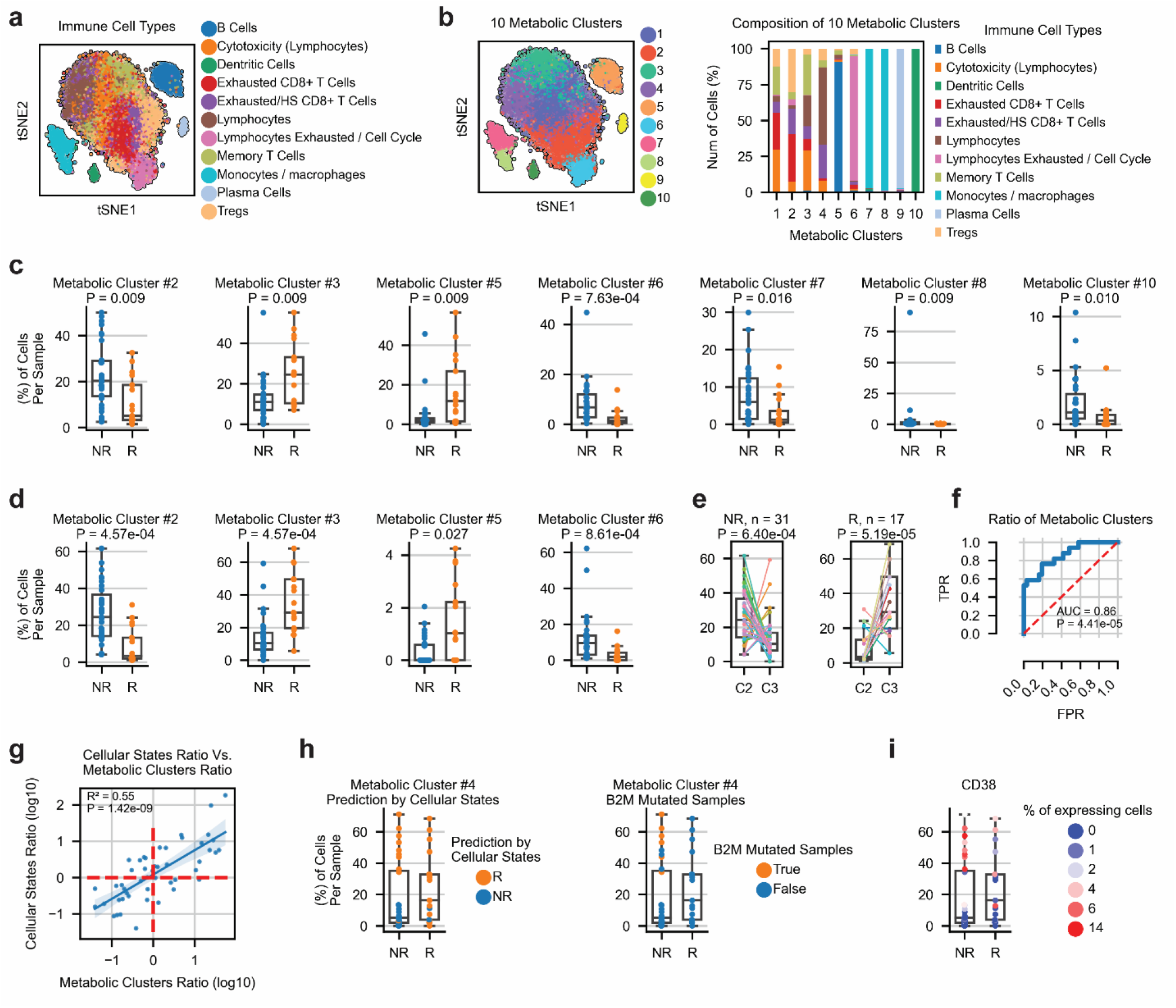
Metabolic-based clusters of melanoma infiltrated immune cells and their association with patient response to ICI. a. Classification of CD45^+^ cells to different types and cellular states according to Sade-Feldman et al.^29^. b. Metabolic clusters of immune cells (left) and their composition by immune cell-types (right). c. Percentage of immune cells found in metabolic clusters associated with patient response, separated by their response status d. Percentage of CD8^+^ T cells found in metabolic clusters associated with patient response, separated by their response status. e. Abundance of CD8^+^ T Cells from metabolic clusters C2 and C3 in each sample by the two response groups. f. The predictive power for response of the metabolic score calculated as the ratio of CD8^+^ T Cells from C3 and C2 in each sample. g. Spearman correlation between the metabolic score of each sample vs. the predictive score suggested by our previous study, Sade-Feldman et al.^29^. h. High abundance of CD8^+^ T Cells from C4 in the misclassified non-responders (left), including B2M mutated samples (right), and their response prediction according to the immunological predictor of Sade-Feldman et al.^29^. i. The fraction of memory and activated CD8+ T cells in C4 out of the total number of CD8^+^ T cells from C4 in each sample, colored according to the expression of CD38.

Unlike these metabolic clusters that corresponded to known immune cell types, each of the metabolic clusters C1-C4 was comprised of a mixture of T cells states, including cytotoxic, exhaustion and memory (Figure 2a, Figure 2b). This finding demonstrates that different immunological states can share metabolic similarity and vice versa. To examine if any of these clusters is associated with patient response, we quantified the fraction of cells in each sample and in each metabolic cluster. We found seven clusters as significantly abundant in one of the response groups (Figure 2c). Two of them, C2 and C3, were among the newly defined metabolic clusters, such that C2 was found to be more abundant in non-responders (two-sided Wilcoxon P-value = 0.009) and C3 was more abundant in responders (two-sided Wilcoxon P-value = 0.009). C2 was enriched with glycolytic genes (PGAM1, PKM, LDHB, GPI), OXPHOS genes from the cytochrome C oxidase and NADH dehydrogenases complexes, as well as antioxidant genes such as SOD1, PRDX2 and PRDX3 (Supplementary Table S3). On the other hand, C3 was characterized by a general down-regulation of metabolic genes without any up-regulated markers (Supplementary Table S3). This finding corresponds to the general downregulation of metabolic pathways seen in cell types associated with responding samples (Figure 1c). At the metabolic pathways level, C2 was enriched with OXPHOS, TCA cycle, glycolysis, and pentose phosphate pathway, while C3 had no significant enrichment of metabolic pathways (Supplementary Table S3). No significant difference between baseline and post-therapy samples was detected in any of the metabolic clusters, in similar to our previous report on the genome-wide clusters^29^. Overall, our semi-supervised analysis detected a metabolic sub-classification of known immunological T cell states, and identified metabolic clusters and genes that were not previously associated with patient response to ICI.

### A metabolic classification of CD8^+^ T cells is predictive of patient response to ICI

Based on the high abundance of CD8^+^ T cells in the clusters associated with patient response, and their role in recognizing tumor antigens, we next focused our analysis on the metabolism of 6350 CD8^+^ T cells found in this dataset^29^. Examining their fraction in the clustering groups C2 and C3, we observed the same direction of association with patient response, but with a higher significance level (two-sided Wilcoxon P-value = 4.57*10^−4^ for both clusters, Figure 2d). Notably, the two metabolic states co-existed in all samples, such that per sample, C2 was generally more abundant in non-responders (two-sided Wilcoxon P-values = 6.4*10^−4^) and C3 was more abundant in responders (two-sided Wilcoxon P-values = 5.19*10^−4^) (Figure 2e). Classifying samples as responders or non-responders based on C3/C2 ratios achieved high predictive power (area under the curve [AUC] of receiver operating characteristic [ROC] = 0.86; two-sided Wilcoxon P-value = 4.41*10^−5^) (Figure 2f).

In our previous study CD8^+^ T cells were similarly clustered into two cellular states, one associated with activation and memory functions and the other with exhaustion^29^. Classifying samples based on the ratio between these immunologic states achieved similar performance with AUC = 0.87. However, the R^2^ between the previous predictor and the current metabolic one is 0.55 (Figure 2g), suggesting that the metabolic score provides additional information. Indeed, a few samples that were misclassified as responders by the immunological predictor due to a high abundance of memory and activated CD8^+^ T cells, were correctly classified using the metabolic predictor as non-responders (Figure 2h, Supplementary Figure S3). This set also includes non-responding samples which had a mutation in the *β*_2_ microglobulin gene, previously used to explain their non-response status (Figure 2h). From a metabolic point of view, we found that nine out of these ten misclassified non-responding samples had a high abundance (30-70%) of CD8^+^ T cells from cluster C4, showing a significant difference between the correctly and incorrectly classified non-responding samples (two-sided Wilcoxon P-value = 4.98*10^−5^, Supplementary Figure S3). While this metabolic cluster was not significantly associated with patient response, we were able to identify several marker genes that significantly differentiate between the true responding samples and the nine samples wrongly classified as responders (Supplementary Table S3). The most significantly up-regulated gene in the samples wrongly classified as responders is CD38 (Figure 2i), a glycoprotein found on the cell surface of many immune cells and converts NAD^+^ to cyclic ADP ribose (cADPR) and to ADP ribose (ADPR)^35,36^. This gene is known to exert immunosuppressive effects through multiple mechanisms, among them is the depletion of NAD^+^ from the tumor microenvironment which impairs T cell activation and differentiation^37^. This finding therefore highlights a sub-cellular state of activated and memory CD8^+^ T cells that expresses CD38 and is abundant in non-responding samples. Overall, our analysis identified a highly predictive metabolic score for classifying patients according to their response status, emphasizing the importance of cellular metabolism in determining patient response.

### Metabolic programs predict response to checkpoint immunotherapy

To generalize these findings into a predictive model that can be used in other datasets, one of two approaches can be taken: in the first, clustering can be performed in each given dataset. However, it would be difficult to receive the exact same clustering solution. The second approach is to use top genes from the two clusters, thus identifying the predictive metabolic cellular states. This solution sets a challenge in this case as the main characterization of C3 was a general down-regulation of metabolic genes. To overcome this obstacle we took a different approach that goes beyond discrete cell clusters and identifies transcriptional metabolic programs that exist as continuous activity programs across and within different cell clusters. This was done by applying consensus non-negative matrix factorization (cNMF)^38^, an approach that performs soft clustering and assigns each cell a gene expression program with a usage value ranging between 0 and 1. We identified ten different metabolic programs (Figure 3a, Methods, Supplementary Table S4), five of them are ‘identity programs’ that are active in specific cell clusters, and the other five are ‘activity programs’ that are active across different cell clusters (Figure 3a). Examining the association of the activity programs with patient response, we found that two of them, P2 and P6, are more abundant in responders and non-responders, respectively (P-value = 0.075 and 5.08*10^−5^, Figure 3b). As expected, these two metabolic programs are also related to the identified C2 and C3 clusters, such that P2 is more abundant in C3 (a two-sided Wilcoxon Rank sums P-value = 4.3*10^−130^) and P6 in C2 (P-value = 8.09*10^−178^, Figure 3e).

**Fig. 3.**
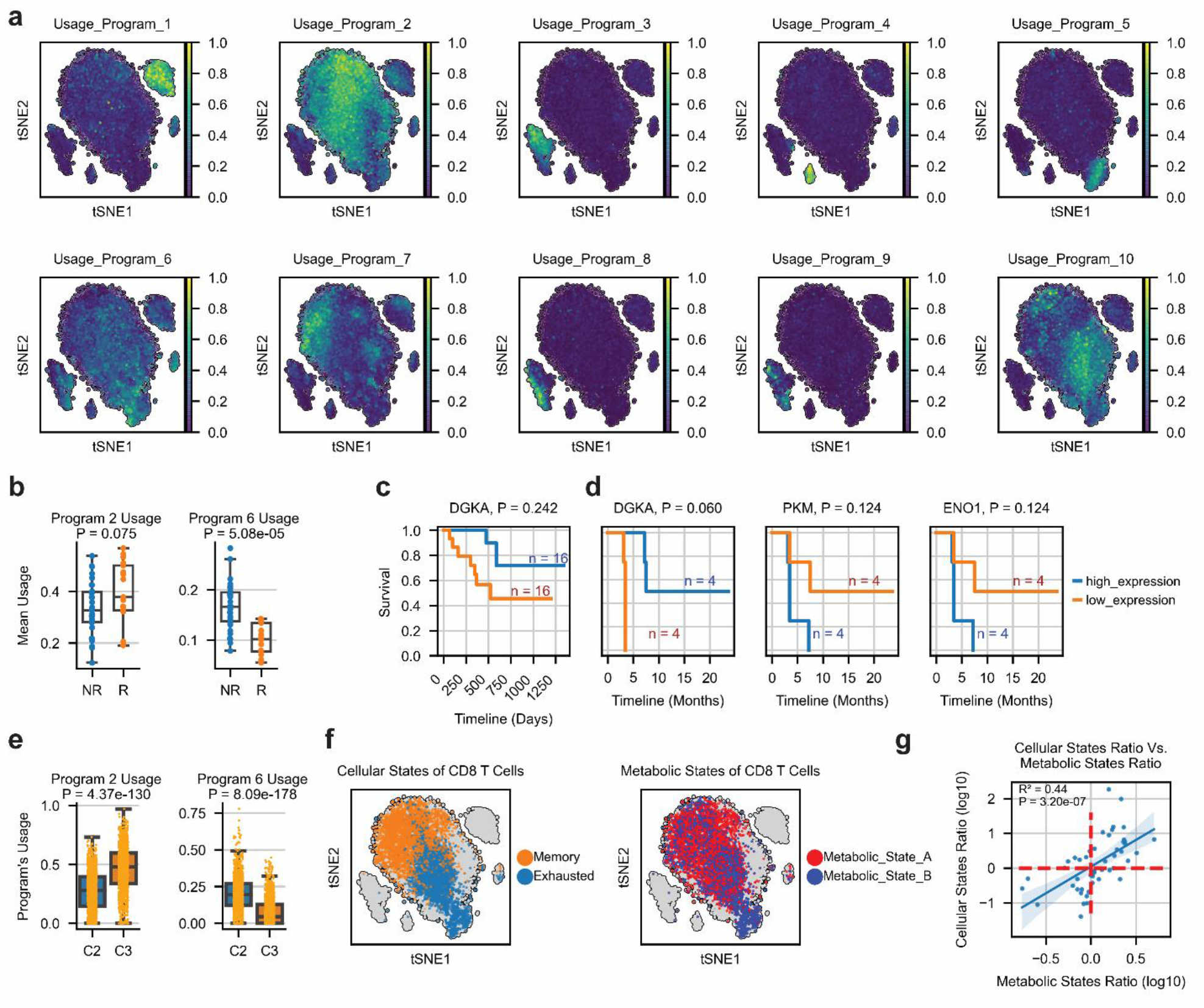
Metabolic programs predict response to ICI. a. 10 metabolic programs obtained using cNMF^38^. b. Distribution of the mean usage values in Programs 2 and 6, separated by the samples response status. c. The association of DGKA (as marker of program P2) with overall survival in melanoma (GSE120575, n=32)^29^. d. The association of DGKA, as well as the glycolytic genes PKM and ENO1 (as markers of program P6) with progression free survival in triple-negative breast cancer (TNBC, n=8) (GSE169246)^51^. e. The usage of programs P2 and P6 in the metabolic clusters C2 and C3. f. Classification of CD8^+^ T Cells into major immunological cellular states, as previously determined^29^ (left) and into major metabolic states (right) according to the top selected markers of each metabolic program. g. Spearman correlation between the metabolic score calculated as the ratio of CD8^+^ T Cells from metabolic states A and B in each sample vs. the predictive score suggested by Sade-Feldman et al.^29^.

To examine the robustness of these programs we analyzed an additional scRNA-seq dataset from non-small cell lung cancer patients (NSCLC) treated with ICI^39^. Similar programs were identified using this dataset, with similar association with patient response (Supplementary Figure S4, Supplementary Table S4). Focusing on these two transcriptional programs, we found that P6 is enriched with OXPHOS

)hypergeometric one-sided P-value = 1.9*10^−10^), Glycolysis (P-value = 3*10^−4^) and Purine synthesis (P-value = 0.005), and P2 is enriched with Inositol phosphate metabolism (P-value = 0.012, Supplementary Table S4, Methods). Indeed, a previous report showed that high OXPHOS in a specific CD8^+^ T cells subset is predictive of immunotherapy resistance in melanoma patients^40^.

Intersecting the gene markers of P2 and P6 between the two datasets, we assembled a narrowed list of the top metabolic markers (Methods, Supplementary Table S4). The first, P2, that we termed ‘Metabolic state A’ contains GSTK1, DGKA, LDHB, MGAT4A, NMRK1, and APRT. The second, P6, that we termed ‘Metabolic state B’ contains PKM, ENO1, COX5A, COX6B1, NDUFB3 and COX8A. While the second group clearly points to activation of glycolysis and OXPHOS, the first contains genes that are active across different metabolic pathways. We however noticed that most exert known protective effect. Specifically, GSTK1 is a glutathione S-transferase that plays a role in cellular detoxification by conjugating reduced glutathione to electrophilic substrates^41,42^. LDHB converts lactate to pyruvate and by that reduces the acidity of the environment^43,44^. DGK deficient mice were shown to contain fewer antigen specific memory CD8^+^ T cells after LCMV infection^45^ and NMRK1 synthesize NAD+, an important factor for T cell activation and differentiation^46^. MGAT4A is an N-acetylglucosaminyltransferase which attaches N-acetylglucosamine (GlcNAc) to the nitrogen atom of an asparagine side-chain, thus playing a crucial role in the regulation of many intracellular and extracellular functions^47,48^. It was also found to be differentially expressed in CD4^+^ and CD8^+^ tumor infiltrating memory T cells in melanoma patients^49^. In hepatocellular carcinoma patients (HCC), it was found to be downregulated in PD1^high^ tumor infiltrating CD8^+^ T cells compared to PD1^intermediate^ CD8^+^ T cells^50^. Here, we found that MGAT4A is differentially expressed in CD8^+^ T cells from samples of responding melanoma patients (Figure 1f). Lastly, APRT is part of the Nucleotide Salvage Pathway that converts Adenine to AMP, and thus may contribute to cell proliferation. Examining the association of these genes with patient survival, we found that the expression of some of them clearly separates between patients, though the difference was not significant after correction for multiple hypotheses due to a relatively small sample size. We found that the expression of DGKA in CD8^+^ T cells is associated with a longer overall survival in melanoma^29^ (adjusted logrank P-value = 0.242, Figure 3c) and a longer progression free survival (PFS) in triple-negative breast cancer (TNBC)^51^ (adjusted logrank P-value = 0.06, Figure 3d). On the other hand, PKM and ENO1 were associated with shorter PFS in triple-negative breast cancer^51^ (adjusted logrank P-value = 0.124 for both genes, Figure 3d).

Based on the expression of these metabolic markers, we divided all CD8^+^ T cells into two major groups of metabolic state A and B (Figure 3f, Methods). We then classified samples as responders or non-responders based on the ratio between the number of cells associated with either of these two metabolic states (Methods). Examining the correlation between this metabolic score and that achieved by our previous immunological one^29^, we found that it explains only 44% of the variance, emphasizing the added information captured by the metabolic classification (R^2^ = 0.44, P-value = 3.2*10^−7^, Figure 3g). We first tested the predictive power of this metabolic score in the above-mentioned melanoma and NSCLC datasets. In melanoma^29^, our predictor achieved an AUC of 0.82 (P-value = 3.05*10^−4^, Figure 4a). A significant difference was maintained also when examining baseline and post-therapy samples separately (P-value = 0.003 and 0.045, respectively). In NSCLC containing only post-therapy samples^39^, our metabolic metric achieved an AUC of 0.8 (P-value = 1.51*10^−4^, Figure 4b). These results were robust to a different number of top markers (Supplementary Figure S5).

**Fig. 4.**
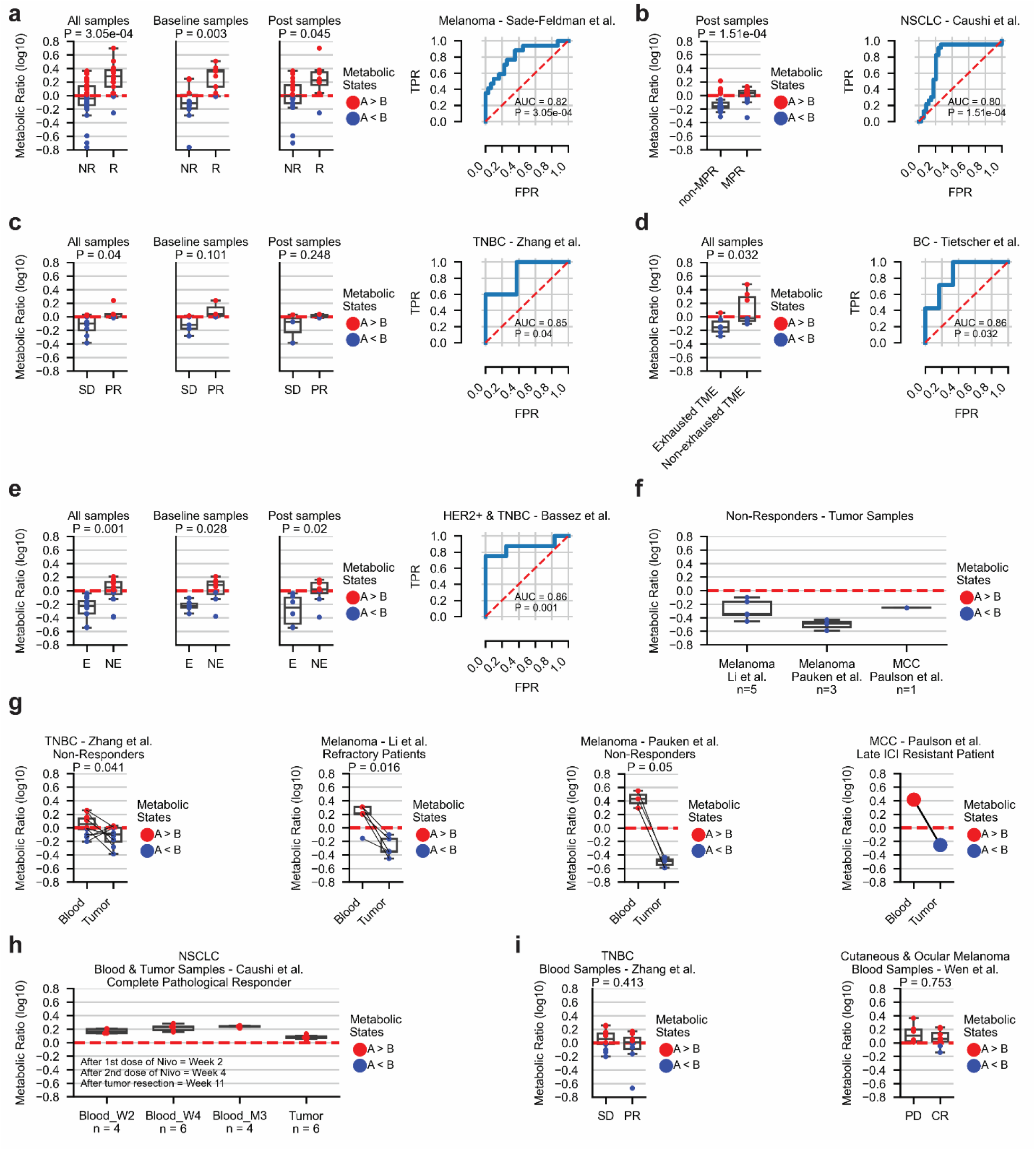
The predictive power of metabolic states across different datasets and cancer types, and its application to blood samples. a-e. The performance of the metabolic predictor in melanoma (a)^29^, NSCLC (b)^39^ and three breast cancer datasets (c-e)^51,54,55^. ROC and the corresponding AUC achieved by the metabolic score are shown on the right; Distribution of the metabolic score in responders and non-responders in the different time points is shown on the left. f. The metabolic states in four datasets including only non-responding patients from three studies, spanning melanoma^40,52^ and MCC^53^. g. A shift in the metabolic states of CD8^+^ T Cells in tumor samples and their matched blood samples taken from non-responding patients with TNBC^51^, melanoma^40,52^, and MCC^53^. h. Similar metabolic states of CD8^+^ T Cells in blood samples and tumor samples of a single complete pathological responder having NSCLC^39^. i. Similar metabolic states of CD8^+^ T Cells in blood samples of responding and non-responding patients having TNBC^51^ and cutaneous and ocular melanoma^57^. Abbreviations: R = Responders, NR = Non-responders, MPR = Major pathological response, SD = Stable disease, PR = Partial response, PD = Progressive disease, CR = Complete response, E = Clonal expansion, NE = No (or low) clonal expansion, TME = Tumor microenvironment.

To further establish the robustness of this predictor, we analyzed seven additional datasets from the public domain having scRNA-seq data from patients treated with ICI^40,51–55^. In a triple-negative breast cancer dataset^51^, our predictor achieved an AUC of 0.85 (P-value = 0.04, Figure 4c). In three melanoma datasets and one Merkel cell carcinoma dataset, all with only non-responding patients^40,52,53^, our predictor correctly classified all samples showing a greater number of CD8^+^ T cells found in metabolic state B (Figure 4f). We also applied our predictor to a dataset of HER2 positive and triple negative breast cancer patients annotated with a clonal expansion phenotype^54^. We found that samples enriched with metabolic state A are more abundant in samples where no (or low) T cell expansion was observed, while metabolic state B was more abundant in samples with a large number of clonal expansion (AUC = 0.86, P-value = 0.001, Figure 4e). This significant trend was observed also when examining baseline and post-therapy samples separately (P-values = 0.028 and 0.02 respectively, Figure 4e). These findings are in agreement with the authors’ original findings that no clonal expansion is enriched in cells with a memory phenotype (TCF7+, GZMK+), while clonal expansion is enriched with cells expressing exhaustion markers (HAVCR2, LAG3, Supplementary Figure S6). They also coincides with a study in melanoma patients showing that highly expanded clonotype families were distributed predominantly in cells with exhausted phenotype having decreased diversity of TCRs^56^.

Finally, we applied our predictor to a dataset containing mostly treatment-naive breast cancer patients having a classification for exhausted and non-exhausted TME provided by the authors^55^. This classification is based on the presence or absence of PD1^high^/CTLA4^high^/CD38^high^ T cells. Our predictor achieved an AUC of 0.86 and significantly differentiated between these two tumor environmental states (P-value = 0.032, Figure 4d). Importantly, our metabolic predictor outperformed the predictor based on immunological cellular states^29^ in all three breast cancer datasets analyzed in this study^51,54,55^, further demonstrating its importance (Supplementary Figure S7).

### The metabolic states of blood-peripheral CD8^+^ T cells is non-predictive of patient response

Applying our metabolic classification to CD8^+^ T cells from blood samples of seven different datasets^39,40,51– 53,57^, spanning melanoma, NSCLC, TNBC, and Merkel Cell Carcinoma (MCC), we found that CD8^+^ T cells in the blood are mostly found in metabolic state A, both in responders and non-responders. This finding corresponds with their general naive-bystander phenotype and lack of checkpoint genes expression. A shift in the metabolic state of CD8^+^ T cells is observed in tumor samples and their matched blood samples taken from non-responding patients with melanoma, TNBC, and MCC^40,51–53^ (Figure 4g). Measuring the metabolic score in blood samples taken at different time points from a NSCLC patient experiencing a complete pathological response^39^, we observed similar ranges of scores as seen in blood samples of non-responding patients (Figure 4h). We also could not identify significant differences between the metabolic scores obtained from blood samples of responding and non-responding TNBC patients^51^, as well as cutaneous and ocular melanoma patients^57^ (Figure 4i). These findings demonstrate that the metabolic state of CD8^+^ T cells is mainly predictive within the TME and emphasize the challenge of identifying reliable blood-based biomarkers.

### A metabolically distinct macrophages cluster in an acquired resistance patient

Clustering the immune cells using the metabolic genes subdivided the original macrophages/monocytes cluster into two metabolic clusters, C7 and C8 (Figure 2b). While all the metabolic clusters were evenly distributed across samples, 82% of the cells associated with C8 were of two specific biopsies (73% and 9%, respectively) from a patient developed acquired resistance following therapy (Figure 5a). Considering known macrophages markers, we found that C7 cells have a higher expression of the RNA editing enzyme APOBEC3A and of interferon related genes such as IFITM1, ISG20, IFI44L & IDO1 (Supplementary Table S3, Supplementary Figure S8), typical to the pro-inflammatory classic macrophages and interferon-primed macrophages^58,59^. C8 cells have a higher expression of metallothionein genes such as MT1F, MT1X, MT1E, MT1G, corresponding to the alternatively activated anti-inflammatory macrophages^60^, together with lipid-related genes (APOE, APOC1, ACP5, FABP5) corresponding to lipid-associated macrophages having canonical M2-like pathways^58,61–63^. Focusing on C8 that was highly enriched in the acquired resistance patient, we found a high expression of several genes in the heme degradation pathway. These include, HMOX1, HMOX2, BLVRA and BLVRB (Supplementary Table S3, Supplementary Figure S8). Interestingly, it was previously shown *in vitro* and in animal models that HMOX1 is involved in the polarization of macrophages into the alternative M2-like phenotype^64^. Here we identify these genes in this unique patient, suggesting a potential mechanism of resistance to ICI, and highlighting metabolic drug targets for enhancing treatment response.

**Fig. 5.**
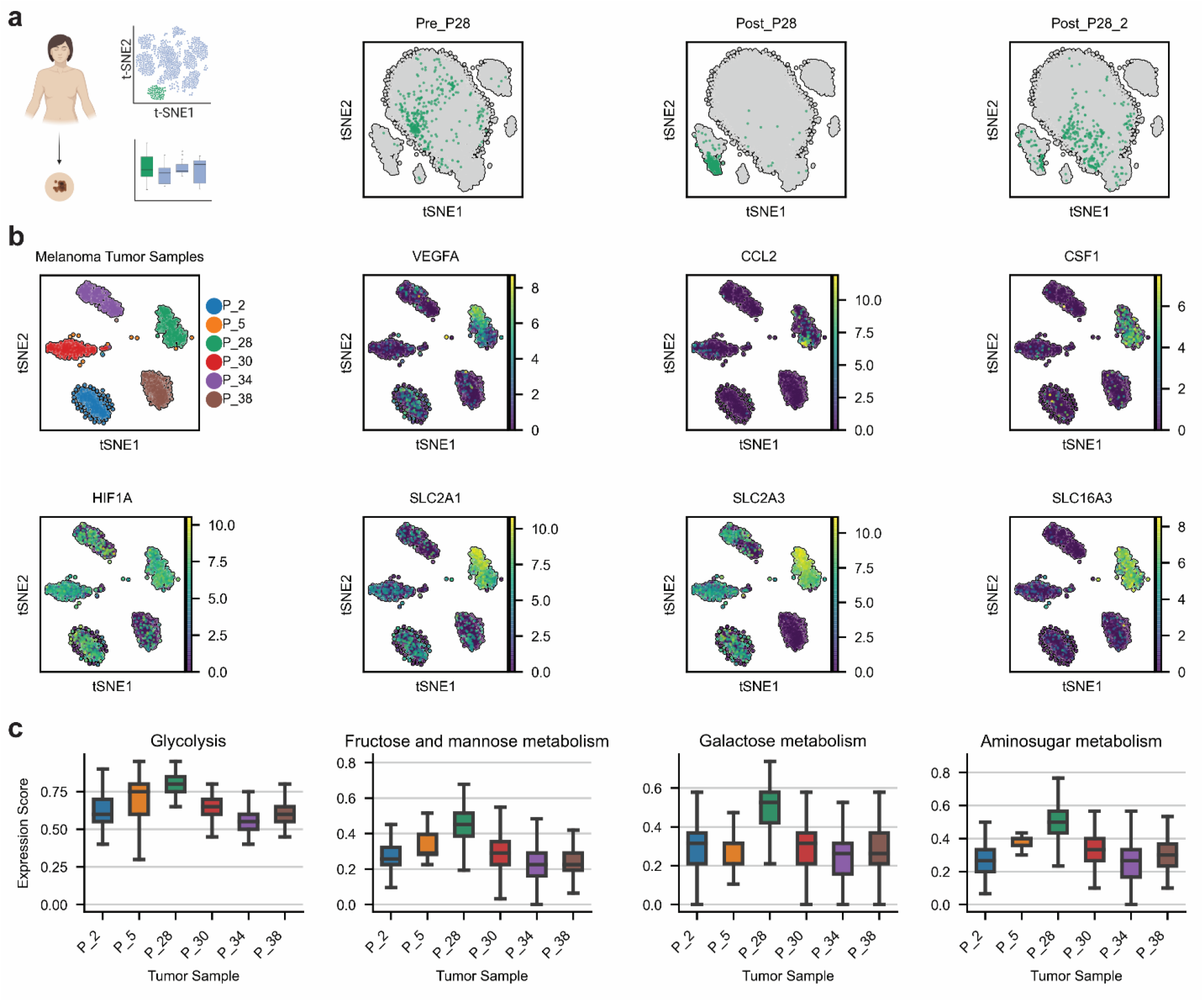
A metabolically distinct macrophages cluster in an acquired resistance patient. a. Three biopsies obtained by a single acquired resistance patient (P28)^29^, with two biopsies containing 73% and 9% respectively of the cells related to metabolic cluster C8. b. Six samples of CD45^-^ malignant cells^65^ and selected differentially expressed genes in the tumor sample of the resistant patient. c. Metabolic score of sugar-related pathways across the different tumor samples.

We next analyzed the corresponding tumor sample of this patient, and compared it to 5 additional tumor samples available in this dataset that lack the M2-like signature^65^ (Figure 5b, Methods). We found a significant increase in the expression of CSF-1, CCL2 and VEGFA (Fisher’s exact test P-values = 4.49*10^−92^, 1.16*10^−29^ and 6.26*10^−68^, respectively), all are tumor-derived factors associated with recruitment of macrophages to the TME and their polarization to the M2-like phenotype^66,67^ (Figure 5b, Supplementary Table S5, Methods). In addition, we identified a significantly higher expression of HIF1A (P-value = 3.15*10^−49^) which is known to induce VEGFA secretion^68^. Considering the metabolic activity of this tumor sample, we found a significantly higher expression of Fructose and mannose metabolism (hypergeometric one-sided P-value = 0.027) and aminosugar metabolism (P-value = 0.027, Figure 5c, Supplementary Table S5, Methods). This tumor also differentially expressed 50% of the genes related to glycolysis and additional genes related to sugar-uptake such as GLUT1 (SLC2A1) and GLUT3 (SLC2A3), compared to a maximum of 2 glycolytic genes in all other tumors (Figure 5b). In addition, we observed a high expression of LDHA in all of these tumor cells, and of MCT4 (SLC16A3), an enzyme catalyzing the secretion of lactate, in 84% of the cells (Figure 5b, Supplementary Table S5). This finding coincides with previous reports regarding the role of lactate in inducing the polarization of macrophages towards the pro-tumorigenic M2-like phenotype, conducted using syngeneic murine tumor models and cancer cell lines^69^. Overall, our findings highlight a specific mechanism of resistance involving a tumor with high activation of sugar-related pathways and its association with anti-inflammatory macrophages, which may be controlled, at least in part, by metabolic genes.

## Discussion

In this study we utilized scRNA-seq data and performed an in-depth transcriptomic characterization of the metabolic changes that occur in immune cells found in the TME of patients treated with ICI. We found a general up-regulation of key metabolic pathways, mostly of oxidative phosphorylation, in cell types associated with non-responding samples such as exhausted cycling cells, dendritic cells and macrophages. On the other hand, we observed a general metabolic down-regulation in cell types associated with responding samples such as B cells and memory T cells. Re-clustering the data based on metabolic genes uncovered novel metabolic clusters associated with patient response, each is a mix of different known immunological cellular states. Furthermore, we detected continuous metabolic programs with different activity levels across CD8^+^ T-cells. Utilizing the top genes of two key programs we were able to accurately classify patients by their response status across various cancer types, including melanoma, Merkel cell carcinoma, lung and breast cancers. Finally, we found that CD8^+^ T cells in blood samples are mostly found in a single metabolic state, regardless of their response status.

An unsupervised analysis of scRNA-seq data typically includes the investigation of all genes passing a set of quality control criteria. This type of analysis naturally highlights the most significant signals found in the data. However, at the same time, it might miss important changes that are weaker and are lost due to multiple hypothesis correction. While this study is a re-analysis of published data, our semi-supervised analysis which was focused on the well-defined biological process of cellular metabolism, was able to uncover significant changes associated with patient response that were not detected in the previous genome-wide analyses done by us and others. We believe that these discoveries were made possible due to the importance of metabolism in immune cells function and interaction with tumor and other cells in the TME.

By identifying metabolic programs with various activity levels, we were able to suggest a sub-classification of CD8^+^ T-cells showing that different immunological states can share the same metabolic state, while similar immunological states can be found under different metabolic states. This new cell classification was predictive of patient response across different cancer types achieving high accuracy levels, that in some cases improved our original classification that was based on the ratio between activation and memory vs. exhausted states^29^. It is therefore plausible that other cellular processes can form additional sub-classifications that will be predictive of patient response.

Metabolism of immune cells has been widely studied, though mainly *in vitro* and in animal models. To the best of our knowledge, this is the first study that metabolically analyzed over 1 million immune cells from human cancer patients treated with ICI, supporting the importance of metabolic processes in determining patient response to therapy. However, in order to fully understand cellular metabolic changes, single-cell proteomic and metabolomic data should be analyzed as well. While these technologies are rapidly advancing, they have not yet reached the scale of transcriptomic data^70–72^. Moreover, functional studies are required to determine the causal effects of metabolic changes on tumor and immune cellular states. Taken together, the increase in clinically annotated data of patients treated with ICI, and the accumulation of additional single-cell omics-data, is expected to significantly advance our understanding of immune cell metabolism and its role in patient response to ICI.

## Methods

### Single-cell RNA-seq datasets and preprocessing

We used 10 scRNA-seq datasets^29,39,40,51–55,57^, with six of them containing tumor biopsies with their matched blood samples, three with only tumor biopsies, and one dataset containing only blood samples (Supplementary Table S1). Expression levels for the single smart-seq2 dataset that we have previously published (GSE120575) were quantified as transcripts per million (TPM)^29^. All other datasets were droplet-based and contained read counts. Expression levels in these cases were first normalized to 10,000 reads per cell, and then log transformed as follows: log_2_(*normalized_counts* + 1). Only samples with defined clinical annotations (response or clonal expansion) were included in the analysis. Furthermore, we considered only patients that were treated with either ICI or a combination of ICI and chemo, as well as a single Merkel cell carcinoma patient that was treated with a combination of T cell therapy and ICI^53^. We also used a dataset of a non-ICI cohort containing mostly treatment-naive patients having annotations for exhausted and-non exhausted TME provided by the authors^55^ (Supplementary Table S1).

### Metabolic genes and metabolic pathways scores

The set of metabolic genes and pathways used in the analysis is based on the Recon 2 metabolic reconstruction^28^. We used the set of 1689 metabolic genes that were expressed in GSE120575 (Supplementary Table S2). The glycolysis pathway as defined in Recon 2 is composed of 78 genes and includes also the gluconeogenesis-related genes. Therefore, to specifically study glycolysis in our analysis, we defined a “Glycolysis” pathway which includes only 20 central genes (supplementary Table S2). To compute a metabolic pathway score for each pathway in every single cell, we calculated for every cell the amount of expressed genes related to each metabolic pathway, out of the total number of genes in that pathway, yielding a ratio between 0 and 1 for each pathway in each cell. Specifically, for each metabolic pathway K, the metabolic pathway score of a single cell was calculated such that:

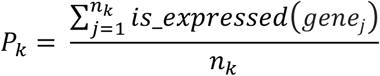

Where *is_expressed*: *x* → {0, 1} and defined as:

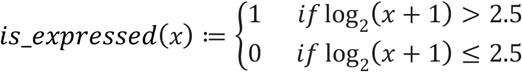

*x* is the expression level in TPM units, *n*_*k*_ is the total number of genes in metabolic pathway K, and the sum is applied on all *j* genes such that *genej* ∈ *pathway*_*k*_

### Metabolic pathway activity and its association with patient response

In order to characterize the metabolic activity of different cell types, we averaged the metabolic pathway scores described above across all cells of each immune cell type. We also ranked the general activity of each metabolic pathway by the mean score of that pathway across all single cells.

To identify metabolic genes that are differentially expressed between responding and non-responding samples, we calculated for every sample the percentage of cells expressing that gene in the two response groups. We included only genes with median expression fraction of 5% across all samples and conducted a two-sided Wilcoxon rank-sums test between the two groups. We then corrected the obtained P-values for multiple hypothesis using the Benjamini-Hochberg false discovery rate^73^. We considered corrected P-values < 0.05 as significant and repeated that process focusing only on CD8^+^ T Cells (Supplementary Table S3). To identify metabolic pathways that are differentially expressed between responding and non-responding samples, we calculated the mean score of each metabolic pathway across all samples and repeated the statistical analysis described above. We conducted this analysis using both all cell types and for CD8^+^ T cells separately (Supplementary Table S3).

### Dimensionality reduction and unsupervised metabolic clustering of immune cells

Principle component analysis (PCA) was applied to all CD45^+^ cells in GSE120575^29^ using 3580 metabolic genes obtained from Recon 3D^74^ (supplementary Table S2). The top 50 components obtained by the PCA were used for the downstream t-distributed stochastic neighbor embedding (tSNE)^75^ with Scanpy’s default perplexity parameter of 30, learning rate of 1,000 and the Euclidean distance metric. The reason we used Recon 3D genes here is that they also contain signaling and regulatory genes, generating a more clear 2D visualization. Importantly, the tSNE algorithm was used only for visualization and was not part of the clustering analysis described below.

To identify cell clusters based on metabolic processes we used Scanpy’s neighbors function with 1689 metabolic enzymes from Recon 2^28^, and generated the K nearest-neighbors (KNN) graph. We then applied the Leiden algorithm^76^ with the default parameters implemented in Scanpy, yielding 10 different metabolic clusters of immune cells (supplementary Table S3). To identify metabolic clusters that are significantly associated with patient response, we calculated for every metabolic cluster the fraction of cells assigned to that cluster in every sample. We then conducted a Wilcoxon rank-sum test between these fractions in responders and non-responders. This process was done for each metabolic cluster and corrected for multiple hypothesis using the Benjamini-Hochberg false discovery rate^73^. Adjusted P-values < 0.05 were considered as significant. We repeated this analysis separately for the set of CD8^+^ T-cells.

### Differential expression analysis

In order to identify differentially expressed genes between two groups *G*_1_ and *G*_2_, we first considered only genes that are expressed (log_2_(*TPM*+ 1) > *thr*) in more than 10% of the cells, in at least one cluster. The threshold (thr) used for smart-seq2 data was ‘2.5’, and ‘1’ for all other 10x datasets. Following, for each gene *i* we count the number of cells in *G*_1_ and *G*_2_ that express it with an expression level (log_2_(*TPM*+ 1) > *thr*) or (log_2_(*TPM*+ 1) **<** *thr*). We then applied Fisher’s Exact test for the corresponding _2_*x*_2_ contingency table. Significant genes were those with an adjusted P-value < 0.05 after correcting for multiple hypothesis using the Benjamini-Hochberg false discovery rate^73^ and log_2_(*FoldChange*) > 0.5. The set of differentially expressed genes in each cluster is listed in supplementary table S3. This analysis was done to identify gene markers in each cluster, comparing it to all other clusters, and for examining differences between responding and non-responding samples.

### Identifying metabolic programs

To identify metabolic programs we applied cNMF^38^ on the raw counts matrix of the melanoma dataset (GSE120575), with the set of 1689 metabolic genes. We filtered out immune cells with total of zero counts and neglected metabolic genes that were expressed in less than three cells. We used 200 NMF replicates for each K and tested the results for K = 5,…,20. Considering the error and stability values for each K, we selected K=13 as the optimal solution. Further reviewing this solution, we filtered out three programs that were active (considering the max usage value) in less than 0.5% of the cells, leaving us with 10 metabolic programs overall. The same process was applied to the NSCLC dataset (GSE176021), obtaining a final set of six programs. Normalized usage values and gene ranks for each program are summarized in supplementary table S4. To identify metabolic programs that significantly differentiate between responding and non-responding samples, we first calculated the mean usage of each program across all CD8^+^ T cells within each sample while excluding programs having median of the mean usage lower than 0.1 across all samples. We performed a two-sided Wilcoxon rank-sums test using the mean usage value of all CD8^+^ T cells within each sample. This process was done for each metabolic program separately and P-values were adjusted using the Benjamini-Hochberg false discovery rate^73^. We considered adjusted P-values < 0.1 as significant for the melanoma dataset, and P-values < 0.15 as significant for the NSCLC dataset. For the melanoma dataset, this process yielded two significant programs (P2 and P6). For the NSCLC dataset, this process resulted with three significant programs (P2, P4 and P6), with the last two having 66% and 33% overlap with the melanoma programs P6 and P2, respectively, considering the top 100 genes in each program.

### Pathway Enrichment analysis

To identify enriched metabolic pathways, we applied a hypergeometric test for each metabolic pathway using the following probability mass function:

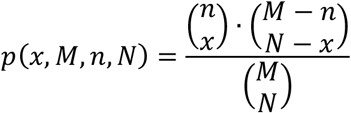

Where *M* is the effective domain size of all of the 1,689 metabolic genes, *N* is the number of genes in the tested list of genes, *n* is the total number of genes associated with each metabolic pathway, and *x* is the number of genes from the tested metabolic pathway found in the tested list of genes. We calculated one-sided P-value for every metabolic pathway using:

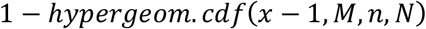

Where *hypergeom. cdf* is the hypergeometric cumulative distribution function implemented by Scipy package version 1.6.2 in Python. We corrected multiple hypothesis using the Benjamini-Hochberg false discovery rate^73^ and considered pathways with adjusted P-value < 0.05 as significant.

This analysis was done using the lists of the differentially expressed genes of metabolic clusters C2, C3, C7 and C8, and is described in supplementary table S3. To identify enriched metabolic pathways in P2 and P6 of the melanoma dataset (GSE120575), we applied this hypergeometric test for each metabolic pathway using the top 100 genes associated with each program, with *N*=100 accordingly. The results of this analysis are summarized in supplementary table S4.

### Identification of CD8^+^ T cells

In our previously published melanoma dataset (GSE120575) we used the original classification of CD8^+^ T cells that was based both on gene markers and manual curation^29^. In all other public datasets we used a stringent criteria such that a CD8^+^ T Cell must express PTPRC (CD45), CD3E and CD8A or CD8B, and cannot express either NCR1, NCAM1 or FOXP3^29^. In all breast cancer datasets of Bassez et al.^54^, Zhang et al.^51^, and Tietscher et al.^55^, we did this process only after focusing on T cells according to the original cell type classification provided by the authors. In the NSCLC dataset of Caushi et al.^39^, we extracted CD8^+^ T cells out of all of the sequenced cells as they went through CD3 positive cell filtration by flow cytometry. In the melanoma datasets of Li et al.^40^, Pauken et al.^52^ and Wen et al.^57^, we extracted CD8^+^ T cells out of all of the sequenced cells as they went through CD8 positive filtration by flow cytometry. Finally, for the Merkel Cell Carcinoma dataset of Paulson et al.^53^, we extracted CD8^+^ T cells out of all unsorted sequenced cells.

### Reconstruction of the metabolic predictor

To reconstruct the metabolic predictor, we considered the two programs that were significantly associated with patient response and had a large overlap between the melanoma and NSCLC datasets^29,39^. In each dataset, genes of the two programs were first ranked according to their usage value. We then calculated the joint rank for the respective programs in both datasets and removed the single gene in each program that corresponded to a housekeeping gene (RPL14 and GAPDH). We eventually selected the top K genes within each program for the reconstruction of the metabolic predictor (Supplementary Table S4). We scored each CD8^+^ T cell based on the number of expressed genes out of the K ‘Metabolic state A’ and the K ‘Metabolic state B’, yielding two scores for each CD8^+^ T cell. Then, each CD8^+^ T cell was classified as A or B based on the majority vote of the two scores: *n*_*state_A_markers*_ ≥ *n*_*state_B_markers*_ for A and *n*_*state_A_markers*_ < *n*_*state_B_markers*_ for B. Finally, we computed a score per sample by taking the ratio between the number of cells classified as belonging to ‘Metabolic state A’ and those belonging to ‘Metabolic state B’. Samples with a ratio > 1 were classified as responders, and those with a ratio < 1 were classified as non-responders. We performed this process on several datasets containing both responding and non-responding samples^29,39,51,54,55^ for different values of K, ranging 3,…,25. For each K we calculated the Receiver Operating Curve (ROC) and the Area Under the Curve (AUC) score, as well as the sensitivity and specificity measures. Best results were achieved for K= 5,6 (Supplementary Figure S5). For simplicity, we selected K=6 for all downstream analysis. A two-sided Wilcoxon rank-sum test was used to calculate the P-value for the performance of the predictor, comparing the metabolic ratio score between true responders and non-responders. For the dataset of Bassez et al.^54^, we conducted this statistical test for the performance of the predictor between samples undergone clonal expansion and those without clonal expansion. For the dataset of Tietscher et al.^55^, we compared between samples defined with exhausted TME and those with non-exhausted TME, provided by the authors.

### Survival analysis

Two datasets in our analysis contained survival data^29,51^. For these two, we performed survival analysis examining whether any of the 12 genes used by our predictor are significantly associated with patient survival. As described above, only the set of CD8^+^ T cells was used in this analysis. For the melanoma dataset^29^, in cases where more than one sample was available for a single patient, we considered only the baseline sample, yielding 32 samples from 32 patients overall. Similarly, for the TNBC dataset^51^ we considered only samples at baseline, yielding 8 samples overall. For each sample we first computed the mean expression of the each gene across all of the associated CD8^+^ T cells. We then split the samples to two groups using the median gene expression value, and conducted log-rank test between the two groups. P-values were corrected for multiple hypothesis using the Benjamini-Hochberg false discovery rate^73^. This process was conducted in each group of six markers (for metabolic states A and B) separately.

### Application of the cellular states predictor

For each one of the three breast datasets utilized in this study^51,54,55^, we calculated for each CD8^+^ T cell identified by the pipeline described above, the fraction of the expressed memory-activated markers and that of the exhaustion markers provided by Sade-Feldman et al.^29^. We then classified each CD8^+^ T cell into one of two cellular states: memory-activated or exhausted, according to the higher fraction of expressed markers. For each sample we calculated the ratio of CD8^+^ T Cells from these two cellular states and examined the predictive power of this ratio, predicting samples as responders if the ratio of memory-activated/exhausted CD8^+^ T cells was higher than 1, and the opposite for classification for negative response (Supplementary Figure S7).

### Melanoma tumor analysis

We utilized seven CD45^-^ samples from 6 patients available in the melanoma dataset^65^. To identify the malignant cells we applied InferCNV (https://github.com/broadinstitute/inferCNV/) and marked those showing copy number variation patterns. We found that all samples had more than 350 malignant cells (out of 384 sequenced cells) except for P5 which had only 24 malignant cells and P28_2 which had only one malignant cell and therefore was combined with the malignant cells from her other available sample. We normalized the TPM matrix as described above and focused on the previously described 1,689 metabolic genes. Differentially expressed genes and enriched metabolic pathways were identified in each tumor according to the pipeline described above (Supplementary Table S5). tSNE^75^ was applied using the set of 1,689 metabolic genes for visualization. For each cell, we scored four sugar-related metabolic pathways as described above: “Glycolysis”, “Fructose and mannose metabolism”, “Aminosugar metabolism”, and “Galactose metabolism”.

## References

1. Pardoll, D. M. The blockade of immune checkpoints in cancer immunotherapy. Nat. Rev. Cancer 12, 252–264 (2012).

2. Fares, C. M., Van Allen, E. M., Drake, C. G., Allison, J. P. & Hu-Lieskovan, S. Mechanisms of Resistance to Immune Checkpoint Blockade: Why Does Checkpoint Inhibitor Immunotherapy Not Work for All Patients? Am. Soc. Clin. Oncol. Educ. B. 147–164 (2019) doi:10.1200/edbk_240837.

3. Gauci, M. L. et al. Long-term survival in patients responding to anti-PD-1/PD-L1 therapy and disease outcome upon treatment discontinuation. Clin. Cancer Res. 25, 946–956 (2019).

4. Prat, A. et al. Immune-related gene expression profiling after PD-1 blockade in non–small cell lung carcinoma, head and neck squamous cell carcinoma, and melanoma. Cancer Res. 77, 3540– 3550 (2017).

5. Hargadon, K. M., Johnson, C. E. & Williams, C. J. Immune checkpoint blockade therapy for cancer: An overview of FDA-approved immune checkpoint inhibitors. Int. Immunopharmacol. 62, 29–39 (2018).

6. Pitt, J. M. et al. Resistance Mechanisms to Immune-Checkpoint Blockade in Cancer: Tumor-Intrinsic and -Extrinsic Factors. Immunity 44, 1255–1269 (2016).

7. Ayers, M. et al. IFN-γ-related mRNA profile predicts clinical response to PD-1 blockade. J. Clin. Invest. 127, 2930–2940 (2017).

8. Riaz, N. et al. Tumor and Microenvironment Evolution during Immunotherapy with Nivolumab. Cell 171, 934–949.e15 (2017).

9. Tumeh, P. C. et al. PD-1 blockade induces responses by inhibiting adaptive immune resistance. Nature 515, 568–571 (2014).

10. Chen, P. L. et al. Analysis of immune signatures in longitudinal tumor samples yields insight into biomarkers of response and mechanisms of resistance to immune checkpoint blockade. Cancer Discov. 6, 827–837 (2016).

11. Daud, A. I. et al. Tumor immune profiling predicts response to anti-PD-1 therapy in human melanoma. J. Clin. Invest. 126, 3447–3452 (2016).

12. Van Allen, E. M. et al. Genomic correlates of response to CTLA-4 blockade in metastatic melanoma. Science (80-.). 350, 207 LP – 211 (2015).

13. Sade-Feldman, M. et al. Resistance to checkpoint blockade therapy through inactivation of antigen presentation. Nat. Commun. 8, p(2017).

14. Roh, W. et al. Integrated molecular analysis of tumor biopsies on sequential CTLA-4 and PD-1 blockade reveals markers of response and resistance. Sci. Transl. Med. 9, eaah3560 (2017).

15. Rizvi, N. A. et al. Mutational landscape determines sensitivity to PD-1 blockade in non–small cell lung cancer. Science (80-.). 348, 124 LP – 128 (2015).

16. Hellmann, M. D. et al. Genomic Features of Response to Combination Immunotherapy in Patients with Advanced Non-Small-Cell Lung Cancer. Cancer Cell 33, 843–852.e4 (2018).

17. Chang, C. H. & Pearce, E. L. Emerging concepts of T cell metabolism as a target of immunotherapy. Nat. Immunol. 17, 364–368 (2016).

18. Renner, K. et al. Metabolic hallmarks of tumor and immune cells in the tumor microenvironment. Front. Immunol. 8, 1–11 (2017).

19. Chang, C. H. et al. Metabolic Competition in the Tumor Microenvironment Is a Driver of Cancer Progression. Cell 162, 1229–1241 (2015).

20. Arner, E. N. & Rathmell, J. C. Metabolic programming and immune suppression in the tumor microenvironment. Cancer Cell 41, 421–433 (2023).

21. Guerra, L., Bonetti, L. & Brenner, D. Metabolic Modulation of Immunity: A New Concept in Cancer Immunotherapy. Cell Rep. 32, 107848 (2020).

22. Fox, C. J., Hammerman, P. S. & Thompson, C. B. Fuel feeds function: Energy metabolism and the T-cell response. Nat. Rev. Immunol. 5, 844–852 (2005).

23. Liu, Y. et al. Tumor-Repopulating Cells Induce PD-1 Expression in CD8+ T Cells by Transferring Kynurenine and AhR Activation. Cancer Cell 33, 480–494.e7 (2018).

24. Watson, M. L. J. et al. Metabolic support of tumour-infiltrating regulatory T cells by lactic acid. Nature 591, 645–651 (2021).

25. Brand, A. et al. LDHA-Associated Lactic Acid Production Blunts Tumor Immunosurveillance by T and NK Cells. Cell Metab. 24, 657–671 (2016).

26. Fischer, K. et al. Inhibitory effect of tumor cell-derived lactic acid on human T cells. Blood 109, 3812–3819 (2007).

27. Elia, I. et al. Tumor cells dictate anti-tumor immune responses by altering pyruvate utilization and succinate signaling in CD8+ T cells. Cell Metab. 34, 1137–1150.e6 (2022).

28. Thiele, I. et al. A community-driven global reconstruction of human metabolism. Nat. Biotechnol. 31, 419–425 (2013).

29. Sade-Feldman, M. et al. Defining T Cell States Associated with Response to Checkpoint Immunotherapy in Melanoma. Cell 175, 998–1013.e20 (2018).

30. Eisenhauer, E. A. et al. New response evaluation criteria in solid tumours: Revised RECIST guideline (version 1.1). Eur. J. Cancer 45, 228–247 (2009).

31. Xiao, Z., Dai, Z. & Locasale, J. W. Metabolic landscape of the tumor microenvironment at single cell resolution. Nat. Commun. 10, 1–12 (2019).

32. Zhang, C. et al. Quantitative profiling of glycerophospholipids during mouse and human macrophage differentiation using targeted mass spectrometry. Sci. Rep. 7, 1–13 (2017).

33. Robichaud, P. P., Boulay, K., Munganyiki, J. É. & Surette, M. E. Fatty acid remodeling in cellular glycerophospholipids following the activation of human T cells. J. Lipid Res. 54, 2665–2677 (2013).

34. Collison, L. W., Murphy, E. J. & Jolly, C. A. Glycerol-3-phosphate acyltransferase-1 regulates murine T-lymphocyte proliferation and cytokine production. Am. J. Physiol. - Cell Physiol. 295, 1543–1549 (2008).

35. Chini, E. CD38 as a Regulator of Cellular NAD: A Novel Potential Pharmacological Target for Metabolic Conditions. Curr. Pharm. Des. 15, 57–63 (2009).

36. Aksoy, P., White, T. A., Thompson, M. & Chini, E. N. Regulation of intracellular levels of NAD: A novel role for CD38. Biochem. Biophys. Res. Commun. 345, 1386–1392 (2006).

37. Kar, A., Mehrotra, S. & Chatterjee, S. CD38: T Cell Immuno-Metabolic Modulator. 1–20 (2020).

38. Kotliar, D. et al. Identifying gene expression programs of cell-type identity and cellular activity with single-cell RNA-Seq. Elife 8, 1–26 (2019).

39. Caushi, J. X. et al. Transcriptional programs of neoantigen-specific TIL in anti-PD-1-treated lung cancers. Nature vol. 596 (Springer US, 2021).

40. Li, C. et al. A high OXPHOS CD8 T cell subset is predictive of immunotherapy resistance in melanoma patients. J. Exp. Med. 219, (2021).

41. Singh, R. R. & Reindl, K. M. Glutathione S-Transferases in Cancer. (2021).

42. Chatterjee, A. & Gupta, S. The multifaceted role of glutathione S-transferases in cancer. Cancer Lett. 433, 33–42 (2018).

43. Wang, Z. H., Peng, W. B., Zhang, P., Yang, X. P. & Zhou, Q. Lactate in the tumour microenvironment: From immune modulation to therapy. EBioMedicine 73, 103627 (2021).

44. Decking, S. M. et al. LDHB Overexpression Can Partially Overcome T Cell Inhibition by Lactic Acid. Int. J. Mol. Sci. 23, (2022).

45. Shin, J., O’Brien, T. F., Grayson, J. M. & Zhong, X.-P. Differential Regulation of Primary and Memory CD8 T Cell Immune Responses by Diacylglycerol Kinases. J. Immunol. 188, 2111–2117 (2012).

46. Ratajczak, J. et al. NRK1 controls nicotinamide mononucleotide and nicotinamide riboside metabolism in mammalian cells. Nat. Commun. 7, 1–12 (2016).

47. Reily, C., Stewart, T. J., Renfrow, M. B. & Novak, J. Glycosylation in health and disease. Nat. Rev. Nephrol. 15, 346–366 (2019).

48. Lau, K. S. et al. Complex N-Glycan Number and Degree of Branching Cooperate to Regulate Cell Proliferation and Differentiation. Cell 129, 123–134 (2007).

49. Li, H. et al. Dysfunctional CD8 T Cells Form a Proliferative, Dynamically Regulated Compartment within Human Melanoma. Cell 176, 775–789.e18 (2019).

50. Kim, H. D. et al. Association Between Expression Level of PD1 by Tumor-Infiltrating CD8+ T Cells and Features of Hepatocellular Carcinoma. Gastroenterology 155, 1936–1950.e17 (2018).

51. Zhang, Y. et al. Single-cell analyses reveal key immune cell subsets associated with response to PD-L1 blockade in triple-negative breast cancer. Cancer Cell 39, 1578–1593.e8 (2021).

52. Pauken, K. E. et al. Single-cell analyses identify circulating anti-tumor CD8 T cells and markers for their enrichment. J. Exp. Med. 218, (2021).

53. Paulson, K. G. et al. Acquired cancer resistance to combination immunotherapy from transcriptional loss of class I HLA. Nat. Commun. 9, (2018).

54. Bassez, A. et al. A single-cell map of intratumoral changes during anti-PD1 treatment of patients with breast cancer. Nature Medicine vol. 27 (Springer US, 2021).

55. Tietscher, S. et al. A comprehensive single-cell map of T cell exhaustion-associated immune environments in human breast cancer. Nat. Commun. 14, (2023).

56. Oliveira, G. et al. Phenotype, specificity and avidity of antitumour CD8+ T cells in melanoma. Nature 596, 119–125 (2021).

57. Wen, T. et al. NKG7 Is a T-cell-Intrinsic Therapeutic Target for Improving Antitumor Cytotoxicity and Cancer Immunotherapy. Cancer Immunol. Res. 10, 162–181 (2022).

58. Ma, R. Y., Black, A. & Qian, B. Z. Macrophage diversity in cancer revisited in the era of single-cell omics. Trends Immunol. 43, 546–563 (2022).

59. Alqassim, E. Y. et al. RNA editing enzyme APOBEC3A promotes pro-inflammatory M1 macrophage polarization. Commun. Biol. 4, 1–11 (2021).

60. Chowdhury, D. et al. Metallothionein 3 Controls the Phenotype and Metabolic Programming of Alternatively Activated Macrophages. Cell Rep. 27, 3873–3886.e7 (2019).

61. Zheng, P. et al. Tumor-associated macrophages-derived exosomes promote the migration of gastric cancer cells by transfer of functional Apolipoprotein e. Cell Death Dis. 9, p(2018).

62. Baitsch, D. et al. Apolipoprotein e induces antiinflammatory phenotype in macrophages. Arterioscler. Thromb. Vasc. Biol. 31, 1160–1168 (2011).

63. Hao, X. et al. Inhibition of APOC1 promotes the transformation of M2 into M1 macrophages via the ferroptosis pathway and enhances anti-PD1 immunotherapy in hepatocellular carcinoma based on single-cell RNA sequencing. Redox Biol. 56, 102463 (2022).

64. Naito, Y., Takagi, T. & Higashimura, Y. Heme oxygenase-1 and anti-inflammatory M2 macrophages. Arch. Biochem. Biophys. 564, 83–88 (2014).

65. Mehta, A. et al. 150 Circulatory plasma proteomic biomarkers predict response to immunotherapy in melanoma patients and reveal biological insights into the tumor microenvironment. A163–A163 (2022) doi:10.1136/jitc-2022-sitc2022.0150.

66. Lin, Y., Xu, J. & Lan, H. Tumor-associated macrophages in tumor metastasis: Biological roles and clinical therapeutic applications. J. Hematol. Oncol. 12, 1–16 (2019).

67. Zhu, S., Yi, M., Wu, Y., Dong, B. & Wu, K. Roles of tumor-associated macrophages in tumor progression: implications on therapeutic strategies. Exp. Hematol. Oncol. 10, 1–17 (2021).

68. Hashimoto, T. & Shibasaki, F. Hypoxia-Inducible Factor as an Angiogenic Master Switch. Front. Pediatr. 3, 1–15 (2015).

69. Colegio, O. R. et al. Functional polarization of tumour-associated macrophages by tumour-derived lactic acid. Nature 513, 559–563 (2014).

70. Lanekoff, I., Sharma, V. V. & Marques, C. Single-cell metabolomics: where are we and where are we going? Curr. Opin. Biotechnol. 75, 1–7 (2022).

71. Seydel, C. Single-cell metabolomics hits its stride. Nat. Methods 18, 1452–1456 (2021).

72. Bennett, H. M., Stephenson, W., Rose, C. M. & Darmanis, S. Single-cell proteomics enabled by next-generation sequencing or mass spectrometry. Nat. Methods 20, 363–374 (2023).

73. Hochberg, Y. & Yosef, B. Controlling the False Discovery Rate : A Practical and Powerful Approach to Multiple Testing. J. R. Stat. Soc.. Ser. B 57, 289–300 (1995).

74. Brunk, E. et al. Recon3D: A resource enabling a three-dimensional view of gene variation in Human metabolism. Nat. Biotechnol. 36, 272–281 (2018).

75. Laurens van der Maaten and Geoffrey Hinton. Visualizing Data using t-SNE. J. Mach. Learn. Res. 9, 2579–2605 (2008).

76. Traag, V. A., Waltman, L. & van Eck, N. J. From Louvain to Leiden: guaranteeing well-connected communities. Sci. Rep. 9, 1–12 (2019).

