## Supplementary Figures for "Metabolic predictors of response to immune checkpoint blockade therapy"

**Supplementary Figures – Metabolic predictors of response to immune checkpoint blockade therapy**

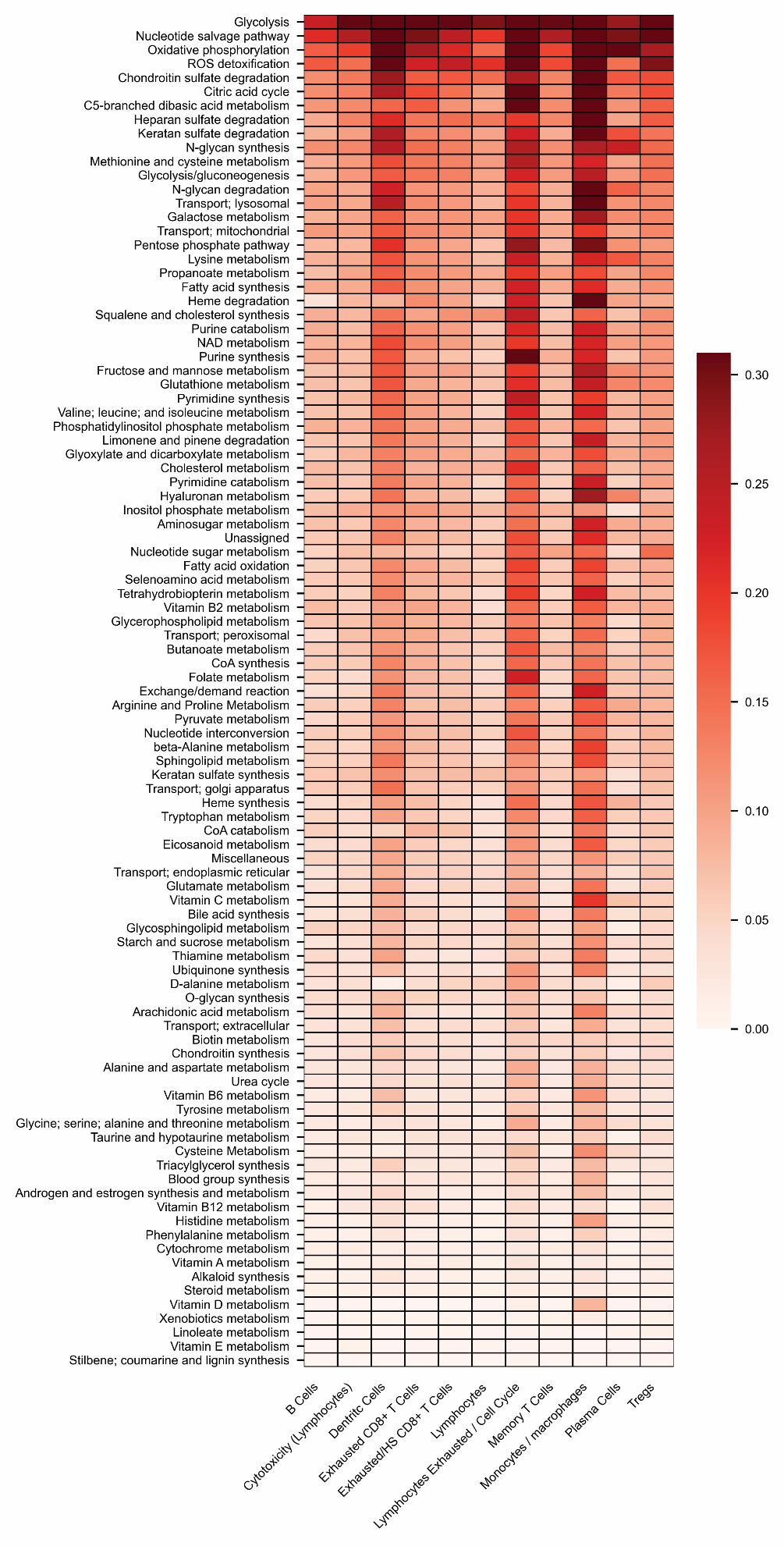

**Supplementary Fig. S1** Ranking of 97 metabolic pathways by their mean score across all single cells in melanoma^1^, and their score across different immune cellular states.

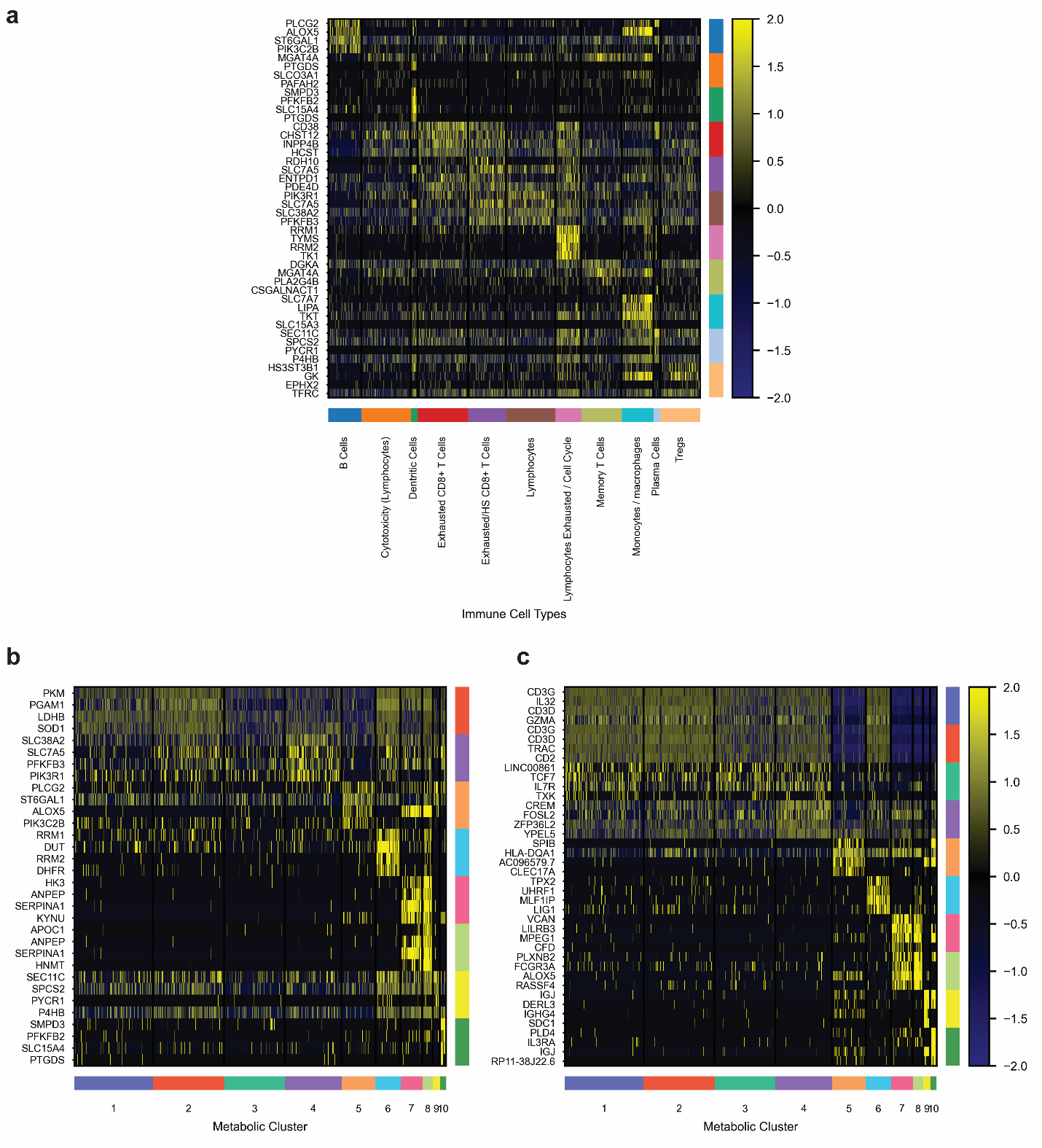

**Supplementary Fig. S2** Differential expression in the Melanoma dataset^1^: a. A heatmap showing differential expression of metabolic genes across different immune cellular states. b. A heatmap showing differential expression of metabolic genes across the 10 metabolic clusters. c. A heatmap showing differential expression of all sequenced genes across the 10 metabolic clusters.

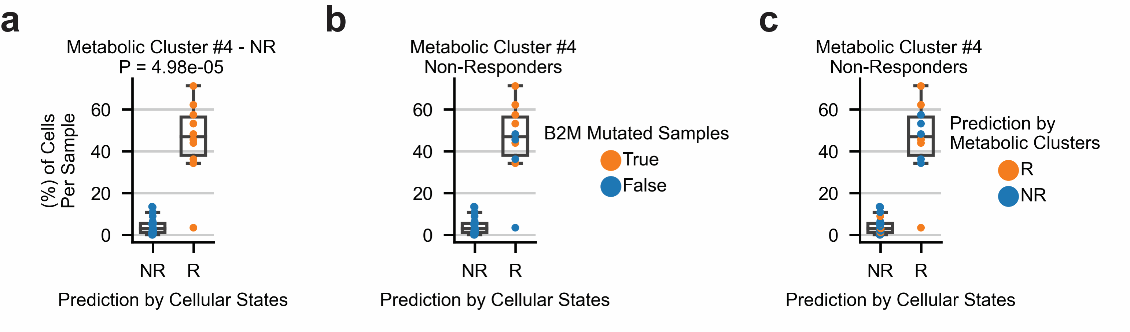

**Supplementary Fig. S3** a. The percentage of CD8^+^ T cells related to metabolic cluster C4 in non-responding samples and their predicted response status by the cellular states predictor suggested by Sade-Feldman et al.^1^ b. $\beta_{2}$ microglobulin mutated samples and their amount of CD8^+^ T cells related to C4, as well as their false prediction as responders according to the cellular states predictor suggested by Sade-Feldman et al.^1^ c. Improved prediction of the misclassified non-responding samples according to the predictor based on metabolic clusters.

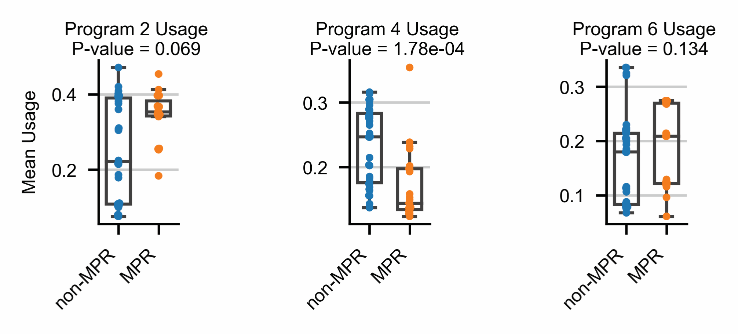

**Supplementary Fig. S4** Distribution of the mean usage values in Programs 2, 4 and 6, separated to samples with MPR and non-MPR in NSCLC^2^.

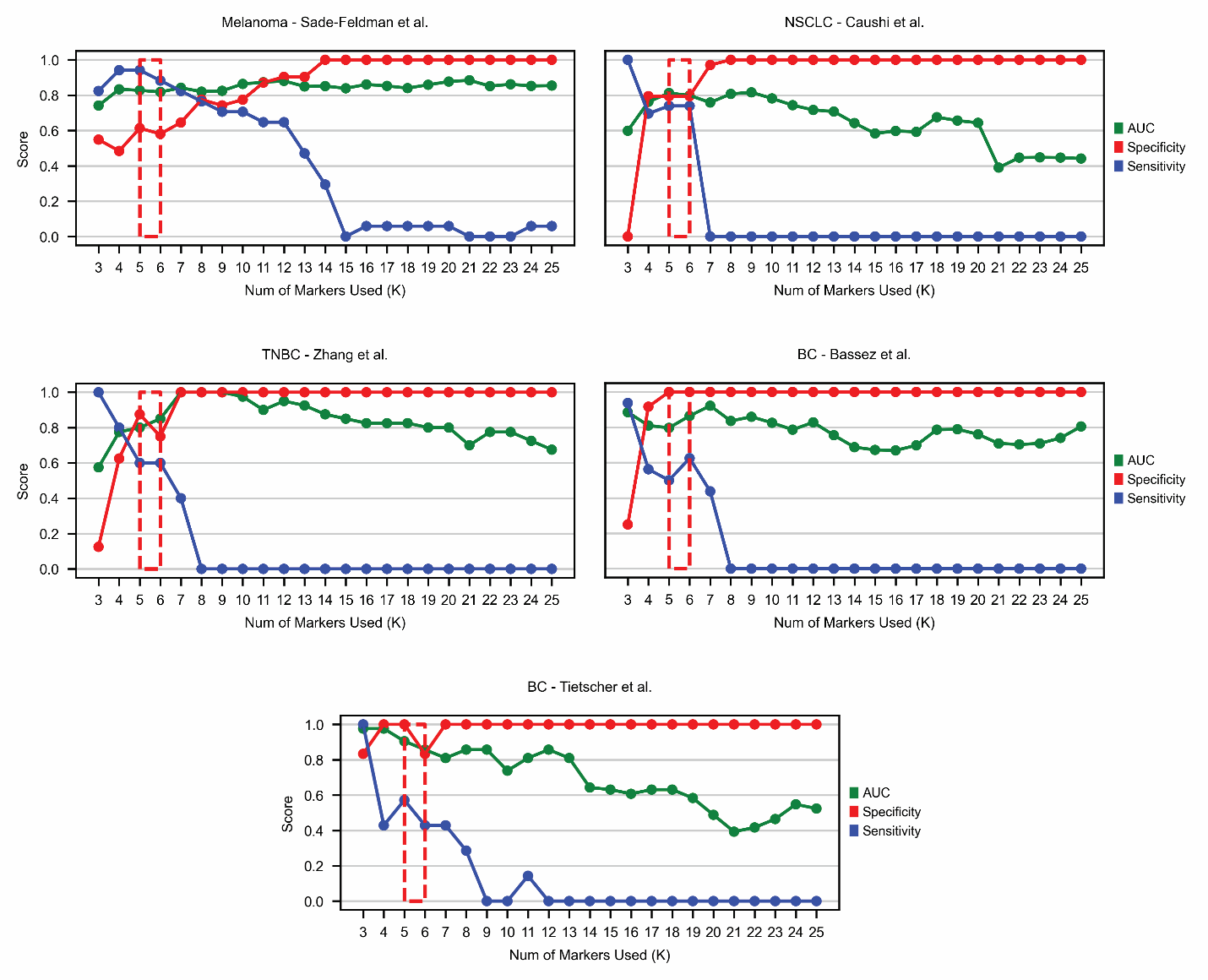

**Supplementary Fig. S5** Robustness of the predictor based on metabolic states of CD8^+^ T cells in melanoma^1^, NSCLC^2^, and breast cancer datasets^3–5^, where K is the number of top metabolic markers used from each one of ‘Metabolic State A’ and ‘Metabolic State B’.

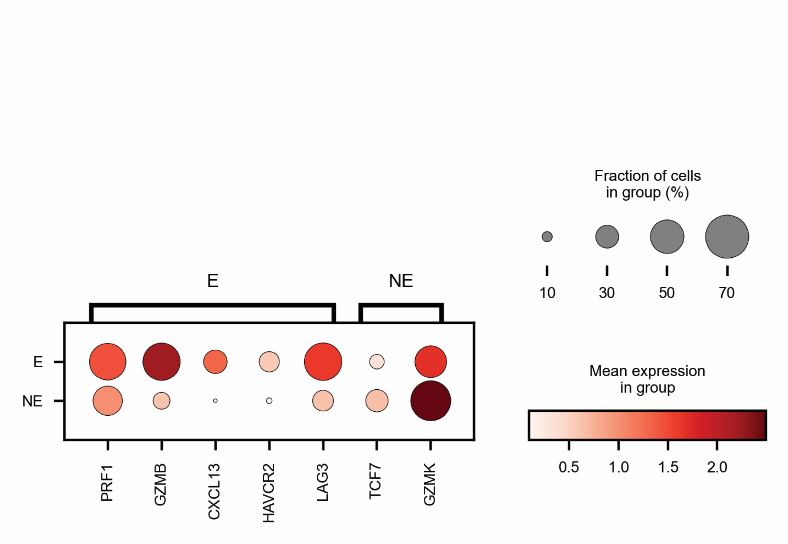

**Supplementary Fig. S6** Expression of memory and exhaustion markers in CD8^+^ T cells by clonal expansion in the breast cancer dataset by Bassez et al.^4^

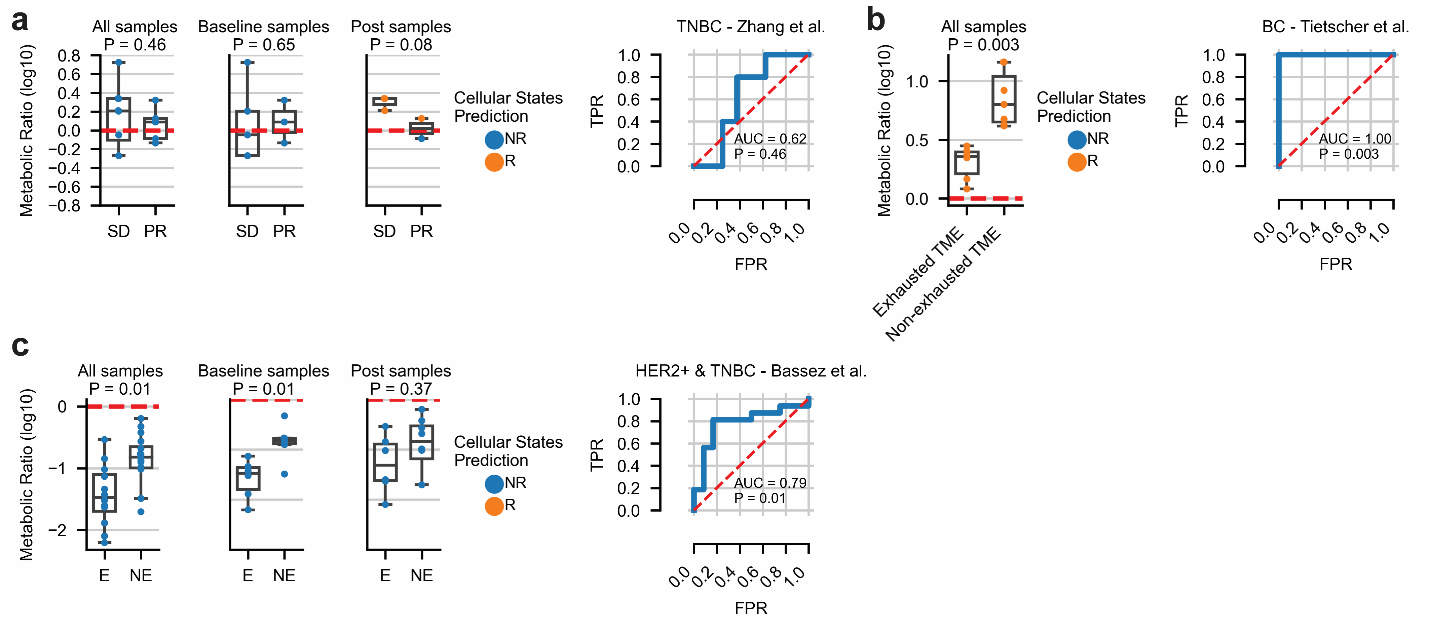

**Supplementary Fig. S7** a-c. The performance of the cellular states predictor suggested by Sade-Feldman et al.^1^ in triple-negative breast cancer (a)^3^, breast cancer (b)^5^, and HER2 positive and triple-negative breast cancer (c)^4^.

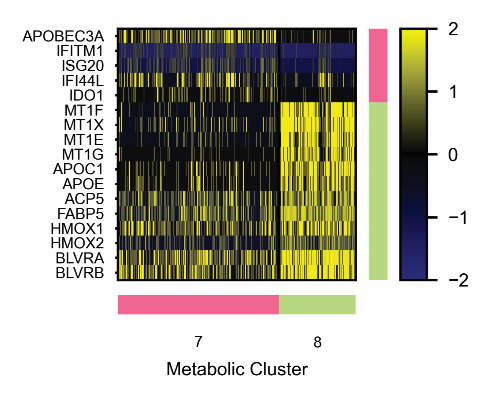

**Supplementary Fig. S8** A heatmap showing selected genes expressed in metabolic clusters C7 and C8, using all sequenced genes^1^.

**References:**

1.	Sade-Feldman, M. *et al.* Defining T Cell States Associated with Response to Checkpoint Immunotherapy in Melanoma. *Cell* **175**, 998-1013.e20 (2018).

2.	Caushi, J. X. *et al.* *Transcriptional programs of neoantigen-specific TIL in anti-PD-1-treated lung cancers*. *Nature* vol. 596 (Springer US, 2021).

3.	Zhang, Y. *et al.* Single-cell analyses reveal key immune cell subsets associated with response to PD-L1 blockade in triple-negative breast cancer. *Cancer Cell* **39**, 1578-1593.e8 (2021).

4.	Bassez, A. *et al.* *A single-cell map of intratumoral changes during anti-PD1 treatment of patients with breast cancer*. *Nature Medicine* vol. 27 (Springer US, 2021).

5.	Tietscher, S. *et al.* A comprehensive single-cell map of T cell exhaustion-associated immune environments in human breast cancer. *Nat. Commun.* **14**, (2023).
